# The fate of sulfamethoxazole and trimethoprim in a micro-aerated anaerobic membrane bioreactor: implications for antibiotic resistance spreading

**DOI:** 10.1101/2023.07.05.547898

**Authors:** Antonella L. Piaggio, Srilekha Mittapalli, David Calderón-Franco, David G. Weissbrodt, Jules B. van Lier, Merle K. de Kreuk, Ralph E.F. Lindeboom

## Abstract

Interest in reusing treated wastewater drives efforts to eliminate antibiotics from water sources to prevent antibiotic resistance. Micro-aerated anaerobic membrane bioreactors (MA-AnMBR) promote wastewater reuse with high organic matter conversion to biogas, under a small footprint. However, the fates of antibiotics, antibiotic-resistant bacteria (ARB), and their antibiotic-resistance genes (ARGs) are not known in these systems. We studied the effects, conversions, and resistance induction, following the addition of 150 μg·L^-1^ of two antibiotics, sulfamethoxazole (SMX) and trimethoprim (TMP), in a laboratory-scale MA-AnMBR. TMP and SMX were removed at 97 and 86%, indicating that micro-aeration did not hamper the removal of the antibiotics. These antibiotics only affected the pH and biogas composition of the process, with a significant change in pH from 7.8 to 7.5, and a decrease in biogas CH_4_ content from 84 to 78%. TMP was rapidly adsorbed onto the sludge and subsequently degraded during the long retention of the solids of 27 days. SMX adsorption was minimal, but the applied hydraulic retention time of 2.6 days was sufficiently long to biodegrade SMX. The levels of three ARGs (*sul1* and *sul2* for SMX, *dfrA1*) and one mobile genetic element biomarker (*intI1*) were analysed by qPCR, in combination with ARB tracked by plating. Additions of the antibiotics increased the relative abundances of all ARGs and *intI1* in the MA-AnMBR sludge, with the *sul2* gene folding 15 times after 310 days of operation. The MA-AnMBR was able to reduce the concentration of ARB in the permeate by 3 log.

**Highlights:**

- Additions of SMX and TMP had a negligible effect on the MA-AnMBR performance.
- The laboratory-scale MA-AnMBR removed 86% of SMX and 97% of TMP.
- A 3 log removal of ARB was achieved between sludge and UF permeate.
- Relative abundances of ARGs were similar in sludge and permeate.
- TMP and SMX resistance is better assessed by the heterotrophic plate count of ARB.

## 1. Introduction

Water demand has been increasing worldwide due to changes in consumption patterns, socioeconomic development, and population growth. Water consumption is expected to rise above one-quarter of the current consumption level by 2050. About 40% of the global population endures water scarcity for one month per year, and 20% lives in countries with high water stress [1]. The use of treated wastewater has risen as a possibility to alleviate water scarcity caused by water stress [2]. Nevertheless, to reclaim treated water, wastewater treatment plants (WWTPs) should be able to provide high-quality effluents. New and upgraded WWTPs should consider not only removing macro-contaminants, such as organic matter, suspended solids, and nutrients but also pathogens and micropollutants [3]. Most conventional WWTPs are not designed for the removal of antibiotics [4, 5] and only minimal removal of pharmaceutical compounds can be observed in the primary treatment of wastewater (i.e., by coagulation, flocculation, and sedimentation) [6]. Currently, several high-income countries are adopting regulations for the treatment of micropollutants and the extension of WWTPs with physical-chemical processes for their removal [7]. However, affordable, and implementable treatment processes with less energy and resource footprints are needed for other regions, such as potentially anaerobic membrane bioreactors (AnMBR).

Antibiotics are important components of human and veterinary medicines. Their consumption is increasing daily, leading to their occurrence in residual waters, such as municipal wastewater and urban and rural run-off. As much as 90% of the consumed antibiotics are excreted without any change in composition or functionality [8]. Among the available antibiotics, sulfamethoxazole (SMX) and trimethoprim (TMP) are found in significantly high concentrations all over the world, as these are some of the most commonly used antibiotics in human and veterinary medications [9]. TMP and SMX are frequently administered together for urinary tract infections [10]. Surface water concentrations of SMX and TMP reached values of 49.7 µg·L^-1^ in Kenia, and 610 µg·L^-1^ in Ecuador [11]. Sim, et al. [12] found high concentrations of SMX and TMP in WWTPs treating wastewater from the pharmaceutical industry, with maximum values reaching 309 and 162 µg·L^-1^, respectively. India’s production of antibiotics is amongst the five top countries worldwide. The Isakavagu-Nakkavagu stream in India carries one of the highest antibiotic concentrations in Asia, with a TMP concentration of 4 μg·L^-1^ [13]. In the water of the largest Barapullah drain in Delhi, the average concentration of TMP was 0.25 µg·L^-1^ [14].

Antibiotics can be removed or transformed by either biotic (biodegradation) or abiotic (sorption, ion exchange, complex formation with metal ions, and polar hydrophilic interactions) processes [15, 16]. On the majority of WWTPs, the sorption and biodegradation of antibiotics occur in parallel. Pharmaceuticals can be biodegraded under aerobic, anoxic or anaerobic conditions, or in combination of all conditions, depending on the antibiotic. The centrally positioned amide group in SMX prevents its degradation under aerobic conditions. However, under anaerobic conditions it can be degraded by reductive cleavage of the molecule due to the adjacently located strong electron-withdrawing sulfonyl group. In the case of TMP, the substituted pyrimidine group can be readily biotransformed under anaerobic conditions [17]. Furthermore, the sorption potential of antibiotics is highly dependent on their molecular charge, polarity, and hydrophobicity, among other characteristics. Hydrophobic antibiotics have a great affinity to solid particles and therefore, have higher chances of being sorbed to sludge particles, o reside long in the process, and to get degraded. The sorption capacity of antibiotics can be described as low, medium, or high, depending on their octanol-water partition coefficient (K_OW_). High sorption is linked to high log K_OW_ values above 4, while low sorption can be considered for antibiotics with low log K_OW_ values below 0.25 [18]. SMX and TMP properties are presented in Table 1.

**Table 1.**
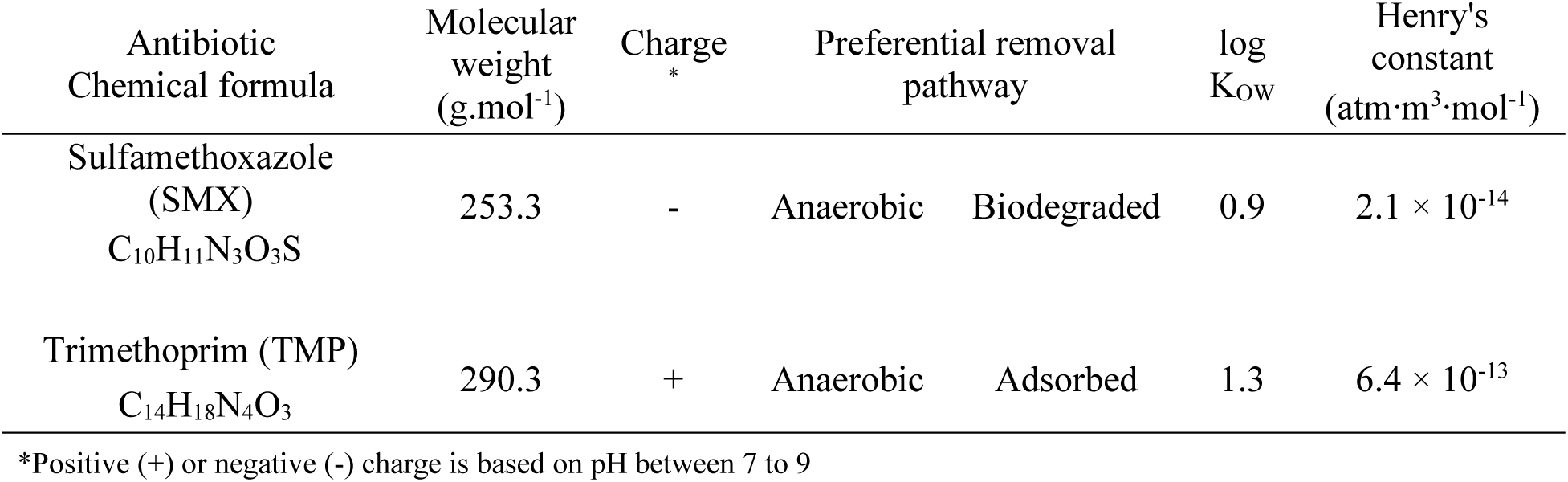
Selected antibiotics and their characteristics. Adapted from NCBI [19] and Alvarino, et al. [17]. K_OW_ is the octanol-water partition coefficient and refers to the sorption capacity of antibiotics. The antibiotics removal pathways refer to the preferential ones, being either aerobic or anaerobic, and biodegraded or adsorbed.

| Antibiotic<br>Chemical formula | Molecular<br>weight<br>(g.mol<sup>-1</sup>) | Charge<br>* | Preferential removal<br>pathway | | log<br>$K_{ow}$ | Henry's<br>constant<br>(atm·m<sup>3</sup>·mol<sup>-1</sup>) |
| --- | --- | --- | --- | --- | --- | --- |
| Sulfamethoxazole<br>(SMX)<br>C<sub>10</sub>H<sub>11</sub>N<sub>3</sub>O<sub>3</sub>S | 253.3 | - | Anaerobic | Biodegraded | 0.9 | $2.1 \times 10^{-14}$ |
| Trimethoprim (TMP)<br>C<sub>14</sub>H<sub>18</sub>N<sub>4</sub>O<sub>3</sub> | 290.3 | + | Anaerobic | Adsorbed | 1.3 | $6.4 \times 10^{-13}$ |

\*Positive (+) or negative (-) charge is based on pH between 7 to 9

The persistence of antibiotics in WWTPs and waterbodies can lead microbial communities to acquire antibiotic resistance. The O’Neil report [20], commissioned by the United Kingdom government, predicts that by 2050, antibiotic resistance infections will lead to 10 million annual deaths, with associated costs above 100 trillion USD. Furthermore, the World Health Organization established that the multi-resistance gained by bacteria is alarming and threatens global public health [21]. Wastewater catchment areas and WWTPs are considered one of the major points of antibiotic resistance release into the environment [22, 23]. Wastewater carries the complex cocktails of chemicals and biological contaminants released from anthropogenic activities. The high density of microbes in WWTPs has been frequently hypothesised to promote the transfer of antibiotic resistance via vertical and horizontal gene transfer in the presence of the antibiotics or other chemicals [24]. The non-resistant bacteria can gain the resistance mechanisms from the antibiotic-resistant bacteria (ARB) via an exchange of mobile genetic elements (MGEs) like plasmids, integrons, and transposons, that contains antibiotic resistance genes (ARGs) [25]. Previous research indicated that the high solids retention times (SRTs) of WWTPs can result in an increased abundance of ARGs in conventional and membrane-based activated sludge systems [26, 27]. However, low convergence in results across the regions has been obtained. The study of more than 60 installations in the Netherlands highlighted that WWTPs mostly abate ARGs of on average 1.8 log gene copies reduction from influent to effluent, i.e., more than 98% removal [28]. Lower removals of ARGs and ARB have been detected under rain events, highlighting the need to improve secondary clarifications under higher hydraulic loadings to remove microorganisms and their ARG pool [29]. Membrane systems can be one solution to enhance the solid-liquid separation.

Membrane bioreactors (MBRs) are potentially effective login treating pharmaceutical wastewater containing various antibiotics and other micropollutants [30]. The SRT and membrane ultrafiltration (UF) pore size are the main operational parameters defining antibiotics removal in MBRs [31]. Several authors found that the optimal SRT to promote antibiotics removal was around 30 days [32–34]. Due to its great affinity to solids, hydrophobic antibiotics can sorb to the sludge particles and then be subjected to biodegradation. The high SRT in MBRs promotes a diverse enzymatic activity due to the manifestation of slowly growing bacteria, which in parallel may support the degradation of the antibiotics [35, 36]. Monsalvo, et al. [37] found that the removal of SMX and TMP in an AnMBR fed with 1.5 μg·L^-1^ of each antibiotic and operated at 30 days SRT was 95.2 and 40%, respectively. Furthermore, Wijekoon, et al. [38] observed a TMP removal of 98% in an AnMBR operated at an SRT of 180 days. Under aerobic conditions, SMX and TMP removal was around 80 and 90% in an MBR system with 70 days SRT [35].

Whether antibiotics are degraded via aerobic, anoxic or anaerobic conversion pathways, determines the need to apply a specific treatment technique, or treatments that combine both redox conditions.

In our previous work, we researched the feasibility of a laboratory-scale micro-aerated (MA) AnMBR, mimicking a full-scale digester equipped with a dissolved air flotation DAF unit for sludge retention instead of a membrane unit (Piaggio et al, 2023, submitted). Results showed improved hydrolysis and negligible effects of the micro-aeration on operation and maintenance of the system. However, thus far, the removal of antibiotics and specifically SMX and TMP in an MA-AnMBR, remains unclear. The application of micro-aeration in an AnMBR might negatively impact their removal efficiency and rate. Since TMP and SMX removal is a mixture of bio-sorption and bio-conversion, the complete solids retention and high SRT provided by the MA-AnMBR system may enhance the degradation of both antibiotics. Furthermore, little is known about the effect of a membrane system on the growth and on the separation of ARB, as well as on the spreading of ARGs, in conjunction with the presence and removal of antibiotics in the wastewater.

Therefore, this study focused on the fate of SMX and TMP, in a laboratory-scale MA-AnMBR and their effect on antibiotic resistance spreading. Antibiotics removal mechanisms (adsorption and/or degradation) and the effects of adding SMX and TMP to the MA-AnMBR feed on its operation and performance were assessed. Measurements of ARGs and ARB from the sludge and permeate of the MA-AnMBR were performed to further understand the complexities and risks linked to the presence of antibiotics in domestic wastewater, and to address the efficiency of MA-AnMBRs to possibly contribute in reducing the spreading of antibiotic resistance from urban water systems.

## 2. Methods

### 2.1. Experimental set-up

A laboratory-scale MA-AnMBR was operated to study the fate of commonly used antibiotics. A continuous-flow stirred tank reactor of 6.5 L liquid volume equipped with an external membrane was used as an anaerobic bioreactor. The membrane system was a side stream inside-out tubular ultrafiltration (UF) membrane, with a pore size of 30 nm (Helyx, Pentair, Minnesota, United States). The process was composed of feed, permeate, aeration, and recirculation pumps (Watson Marlow 120U and 323U, Falmouth, United Kingdom). A Memosens CPS16D combined sensor (Endress+Hauser, Reinach, Switzerland) was used to measure the sludge oxidation-reduction potential (ORP), pH, and temperature in a continuous way. The MA-AnMBR operational conditions are based on the method described by Piaggio et al, 2023 (submitted), and summarized in Table 2. Micro aeration to the system was introduced in the reactors bulk liquid, in three sets of four hours of aeration followed by four hours of no aeration. The total daily air volume introduced to the system was around 120 mL, which corresponds to 25 mL O_2_ (based on oxygen to air ratio of 0.21). The reactor setup and scheme are depicted in Figure 1. The concentrated influent consisted of synthetic wastewater, following an adapted recipe from Ozgun [39]. The synthetic feed had an average COD of 4.9 ± 0.6 g·L^-1^, 66.4 ± 3.4 mgPO ^3+^-P·L^-1^, and 244 ± 8 mgNH_4_^+^-N·L^-1^. Feed composition and its recipe are presented in **Annex A**.

**Figure 1.**
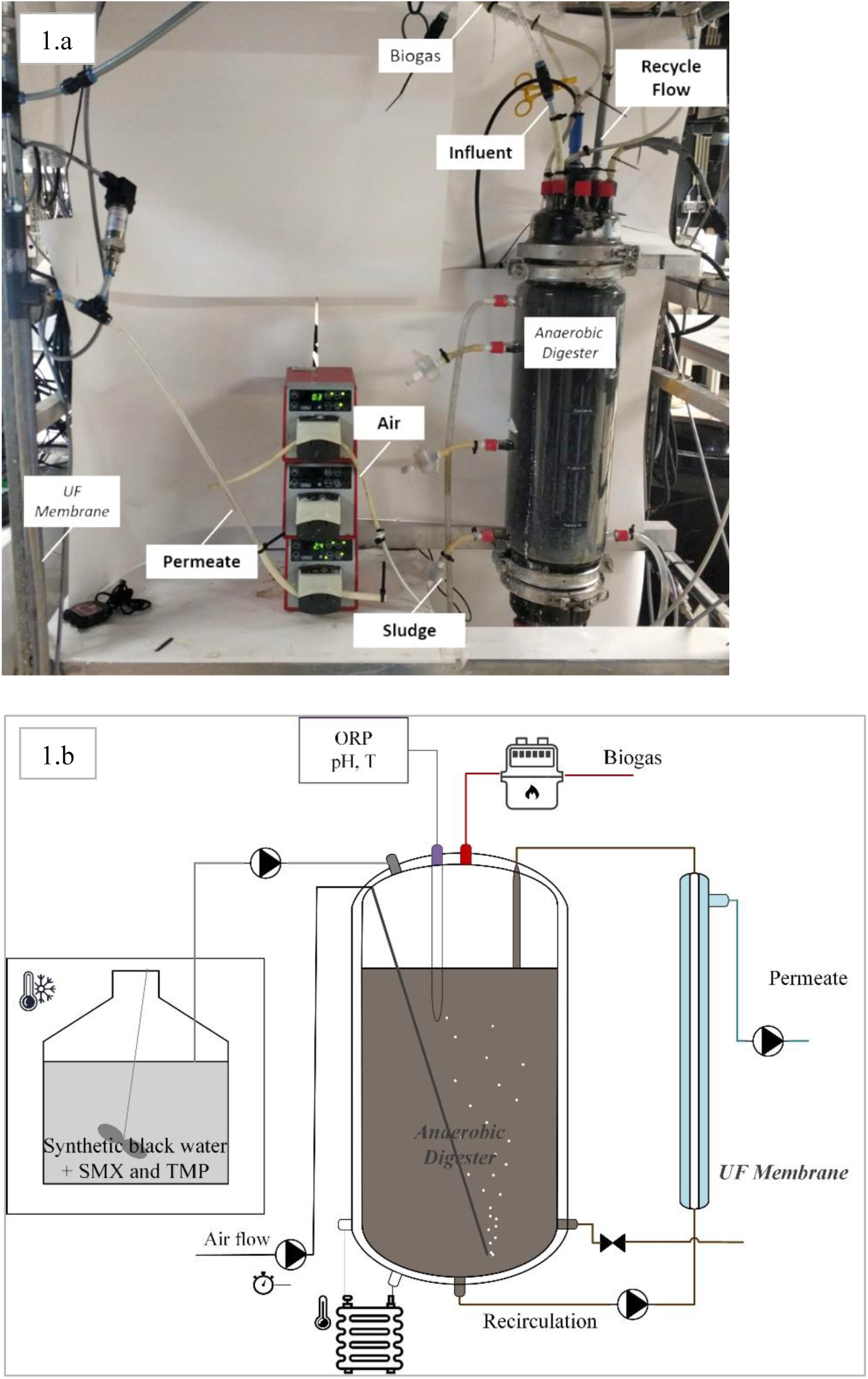
Micro-aerated AnMBR setup (MA-AnMBR). Adapted from Piaggio et al, 2023 (Submitted). Figure 1.a shows the MA-AnMBR lab-scale setup. The figure 1.b is a schematic representation of the lab-scale unit.

**Table 2.** Micro-aerated AnMBR running conditions.

|  | Unit | Value |
| --- | --- | --- |
| Feed flow | L·d<sup>-1</sup> | 2.5 |
| Permeate flow | L·d<sup>-1</sup> | 2.3 |
| Reactor volume | L | 6.5 |
| Temperature | °C | 37 |
| Hydraulic retention time (HRT) | d | 2.6 |
| Solids retention time (SRT) | d | 28 |
| Organic loading rate | gCOD·L<sup>-1</sup>·d<sup>-1</sup> | 1.9 |
| Recirculation flow | L·d<sup>-1</sup> | 1300 |
| Crossflow velocity | m·s<sup>-1</sup> | 0.6 |
| Membrane flux | LMH | 10.0 |

### 2.2. Use of antibiotics SMX and TMP

Removal by adsorption and biodegradation of SMX and TMP was studied in both batch-scale systems and continuously operated MA-AnMBRs. From the literature, SMX and TMP concentrations in influent of WWTPs vary from 10 to 500 μg·L^-1^. Considering that the synthetic influent of the MA-AnMBR is concentrated wastewater, 150 µg·L^-1^ of each antibiotic was added to the feed of the lab-scale MA-AnMBR. The addition of the antibiotics was done in steps and is described below. Moreover, SMX and TMP removal by adsorption was studied in batch tests (described in 2.5 hereafter) with concentrations between 10 and 150 µg·L^-1^.

### 2.3. Reactor phases

The MA-AnMBR system was operated under stable conditions for 90 days, before the addition of antibiotics was started. This phase is referred hereafter to as *P.I*.

The subsequent phase called *P.II* refers to the time frame in which the two antibiotics, TMP and SMX, were added step-wise to the reactor feed, as following. Firstly, TMP was added to the feed in three steps of increasing concentrations: 10, 50, and 150 µg·L^-1^. The time lapse between each concentration shift corresponded to 3 hydraulic retention times (HRT), which was approximately one week. Thereafter, SMX addition started, and was done similarly to TMP in the same concentration steps and time. The whole *P.II* phase lasted 40 days.

Once both the SMX and TMP feed concentrations were 150 µg·L^-1^, the reactor was continuously fed for a period of 120 days with the above-mentioned synthetic concentrated wastewater and 150 µg·L^-1^ antibiotics. This phase is referred to as *P.III*.

Finally, the influent without antibiotics was fed again to the system from day 250 onwards and monitored for a further 180 days, reaching a total operational time of 430 days. This last monitoring phase was denominated *P.IV*. A schematic overview of the reactor phases is displayed in Figure 2.

**Figure 2.**
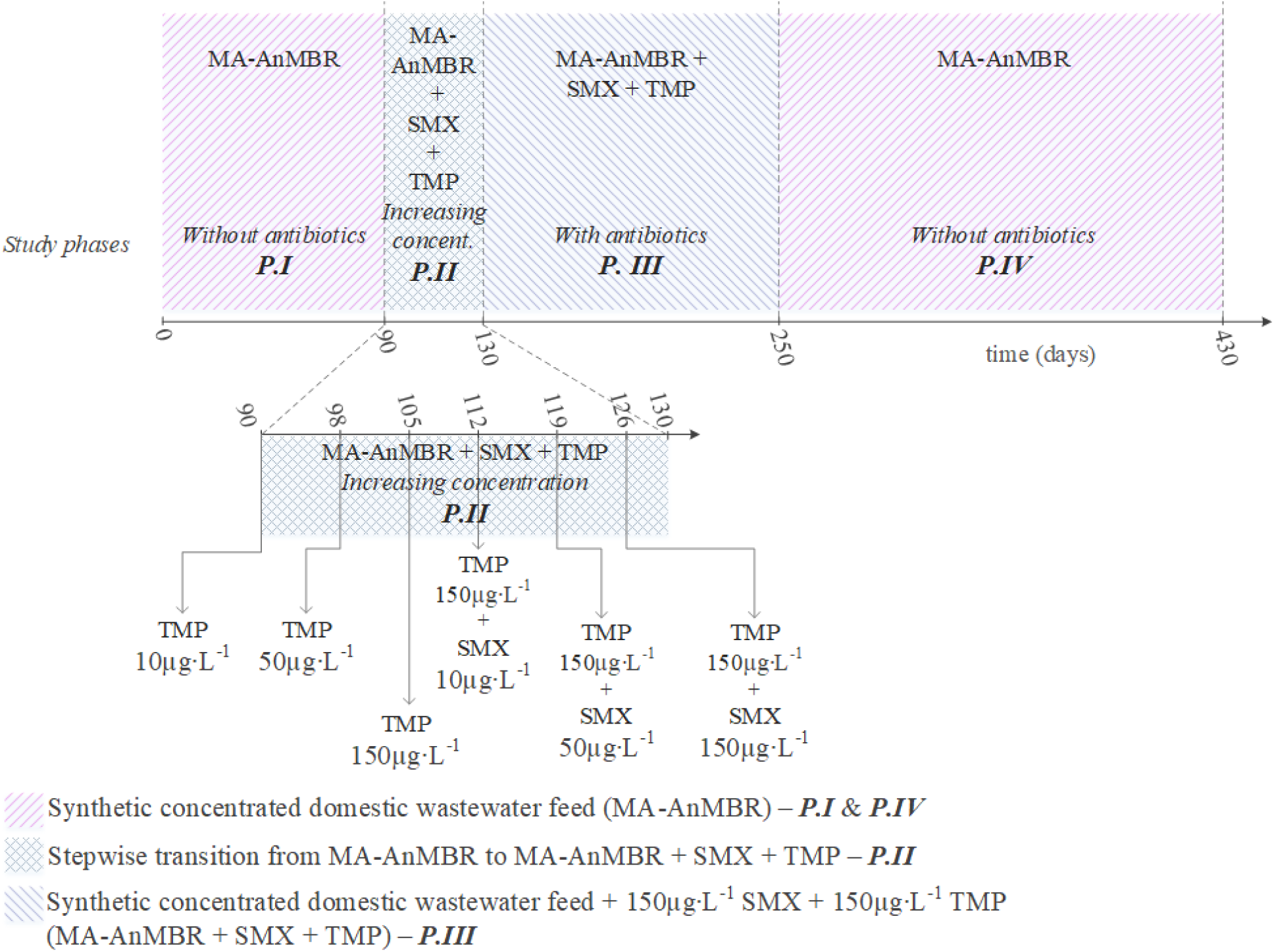
Schematic representation of the reactor phases.

### 2.4. Analytical methods

COD measurements were conducted using HACH Lange test kits LCK 314, 514, and 014 (HACH, Tiel, The Netherlands). The nutrients, orthophosphate (PO_4_^3-^-P), total nitrogen (TN), ammonium-nitrogen (NH_4_^+^-N), and nitrate-nitrogen (NO_3_^-^-N), were measured with HACH Lange test kits (LCK 238, 303, and 339, respectively). Total and volatile solids (in triplicates) were measured according to the APHA-Standard Methods [40].

For the analysis of biogas, the gas samples were collected using 1.5 mL sterile syringes after which they were immediately injected into an Agilent tech 7890A gas chromatograph (GC) (Agilent, Santa Clara, CA, U.S.A) equipped with a thermal conductivity detector. To analyse the composition of the gas samples, two separate columns, HP-PLOT U and a Molesieve GC column (Agilent 19095P-MS6, Santa Clara, CA, U.S.A) of 60 m x 0.53 mm x 200 μm were used. Helium at a flow rate of 10 mL·min^-1^ was used as a carrier gas. The operational temperatures for the injector and detector were 40 °C and 200 °C respectively [41].

The composition of the volatile fatty acids (VFA) of the MA-AnMBR sludge were measured using an Agilent tech 7890A GC, with helium as a carrier gas. The gas flow rate was 2.45 mL·min^-1^ (at 0.76 bar), and detector and injector temperatures were 225 °C and 240 °C, respectively. The liquid samples were collected in 2 mL Eppendorf every week and measured following the procedure described by Garcia Rea et al [42]. Acetic, caproic (IC6), and propionic acids are measured in mg·L^-1^, and the final VFA concentration is expressed in meq·L^-1^ based on the molecular weight of each compound.

#### 2.4.1. Antibiotics concentration measurement

Antibiotics concentrations in samples from batch experiments and the continuous-flow reactor were analysed using liquid chromatography-tandem triple quadrupole mass spectrometry (LC-MS). For feed and permeate samples, before analysis, a volume of 2 mL of sample was centrifuged in a micro-centrifuge (Eppendorf, Hamburg, Germany) at 10,000⊆g for a period of three minutes, and the supernatant was thereafter filtered through a 0.20 µm syringe filter (Chromafil® Xtra PES 20/25, Macherey-Nagel, Germany). Samples were stored at -20°C pending LC-MS analysis.

Antibiotics concentrations in the sludge from the continuous-flow reactor were measured using the methods described by Wijekoon, et al. [38]. Homogenous sludge samples of 15 mL were centrifuged (Sorval ST 16R Centrifuge, Thermo Fisher Scientific, Waltham, MA, U.S.A) at 14,000⊆g for 15 minutes, and the supernatant was discarded. After freezing the remaining sludge samples at -80°C for at least one day, the samples were freeze-dried for 24 hours (BK-FD10, Biobase, Shandong, China). The dried sludge was then grounded to a fine powder using a hand mortar and pestle, and a mass of 0.4 g was transferred to a tube, where 4 mL of methanol (>99%) was added. The samples were mixed with a vortex and sonicated for 10 minutes with an amplitude set up of 20% and temperature less than 60°C (Branson 450 Digital Sonifier, Connecticut, U.S.A). Afterwards, the sample was centrifuged at 3,300⊆g for 15 min, and the supernatant was collected in a fresh 15 mL tube for further analysis. Finally, the solution was filtered through a 0.20 μm syringe filter (Chromafil® Xtra PES 20/25, Macherey-Nagel, Germany), and treated in a similar way to the permeate and liquid samples.

Chromatographic separation of the pharmaceuticals was performed by the ACQUITY UPLC® BEH C18 column (2.1 × 50 mm, 1.7 μm, Waters, Ireland) with a gradient elution of ultrapure water. Acetonitrile was the mobile phase and its flow rate was set to 0.35 ml.min^−1^ using an ACQUITY UPLC I-Class Plus pump (Waters, U.S.A). Ultrapure water and acetonitrile (LC-MS grade, Biosolve, France) were acidified with 0.1% formic acid (LC-MS grade, Biosolve, France). Detection of the pharmaceuticals by mass spectrometry (Xevo TQ-S micro, Waters, U.S.A) was conducted in the positive and negative electrospray ionization modes. The obtained data were analysed and compared to internal standards, based on the methods and information described by Zheng, et al. [43].

#### 2.4.2. Heterotrophic plate count

Microbiological screening and quantification were performed by spread plate method according to APHA-Standard Methods [40]. The total heterotrophic bacteria count was assessed by plating 0.1 mL of sample (either from the permeate or sludge) on a non-selective tryptone soya agar and low-nutrient Reasoner’s 2A (R2A) agar. ARB were measured by adding concentrations of 50.4 mg·L^-1^ of SMX, or 16 mg·L^-1^ of TMP to the plate media (R2A). The antibiotics concentrations added to the R2A media were chosen based on the minimum inhibitory concentrations given by the Clinical and Laboratory Standards Institute [44] and studies performed by Zarei-Baygi, et al. [24], [45].

### 2.5. Adsorption batch tests

Adsorption tests were performed in 250 mL glass bottles at 10°C for a period of seven hours, to inhibit biodegradation. A volume of 100 mL of acclimated sludge from the first phase *P.I*, collected daily and stored at 4 °C, was used to perform all adsorption experiments. For SMX and TMP, three different antibiotic concentrations were tested: 10, 50, and 150 µg·L^-1^. Each antibiotic concentration was added to the 250 mL glass bottles. All experiments were performed in triplicates, summing up to a total of 18 bottles and six different analysis conditions. Immediately after the addition of the antibiotics, the bottles were placed on the magnetic stirrer at 160 RPM (C-MAG HS7, IKA®, Staufen, Germany), for over six hours. Sludge samples were collected in Eppendorf tubes of 2 mL volume and performed every five minutes for a period of 45 minutes, then every 15 minutes for two hours, and finally, every half an hour for the next four hours. After collecting, the samples were immediately centrifuged in a micro-centrifuge (Eppendorf, Hamburg, Germany), the supernatant was filtered through a 0.20 μm syringe filter, and the samples were stored at -20°C until further analysis of the residual dissolved pharmaceuticals in the LC-MS.

### 2.6. Genetic analysis of ARGs

#### 2.6.1. DNA extraction

Triplicate sludge and permeate samples from the MA-AnMBR were taken across the four studied phases *P.I to P.IV* for molecular analyses of their ARG content, after extraction of their total DNA. DNA extractions were performed following the protocol established by Albertsen, et al. [46], with minor adaptations as following.

Homogenised sludge samples of 1.5 mL were transferred into an Eppendorf tube and centrifuged in a micro-centrifuge (Eppendorf, Hamburg, Germany). Approximately 50 mg of sludge pellets were added to DNA extraction tubes of the soil FastDNA spin kit (MP Biomedicals, Irvine, CA, U.S.).

MA-AnMBR permeate volumes of 2.0 L (per sample) of permeate were collected anaerobically and filtered through a 0.20 µm filter (Chromafil® Xtra PES 20/25, Macherey-Nagel, Germany). Afterwards, the filter was cut using a scissors and the pieces were transferred in 1.5 mL Eppendorf tubes with tweezers (all used material was previously sterilized using autoclave (Fedegari FVA3/A1, Albuzzano, Italy). Autoclaved conditions were achieved by pressure-sterilization at 121°C for 20 min.

The concentrations of the DNA extracts were measured using a Qubit dsDNA assay kit (Thermo Fisher, Waltham, MA, U.S.A). Finally, the DNA samples were frozen at -25 °C until they were sent for gene amplification (Novogene Europe, Cambridge, United Kingdom).

#### 2.6.2. Quantitative polymerase chain reaction (qPCR) analysis of selected ARGs

Three ARGs were selected for qPCR analysis on the DNA fractions extracted from the sludge and permeate of the continuous MA-AnMBR lab-scale system. The chosen ARGs targeted the sulphonamide resistance genes *sul1* and *sul2*, and dihydrofolate reductase gene *dfrA1*, to assess the potential resistance gained by the sludge by the addition of SMX and TMP, respectively. Aside from the ARGs, one mobile genetic element (MGE) biomarker was selected to investigate gene mobility, namely the class I integron-integrase gene *intI1* [47]. The 16S rRNA gene was selected as a proxy to quantify total bacteria. Standards, primers, and mix solutions were based on the work performed by Calderón-Franco, et al. [48], and are given in **Annex B** of the supplementary material.

## 3. Results

### 3.1. MA-AnMBR performance

Changes in the MA-AnMBR performance before and after adding the antibiotics are shown in Table 3. The COD removal was always above 97%, with permeate COD values that varied between 50 and 90 mg·L^-1^. The statistical difference between the reactor parameters was assessed using ANOVA single-factor between the MA-AnMBR parameters before and during the antibiotic’s addition phases (*P.I and P.III*).

**Table 3.**
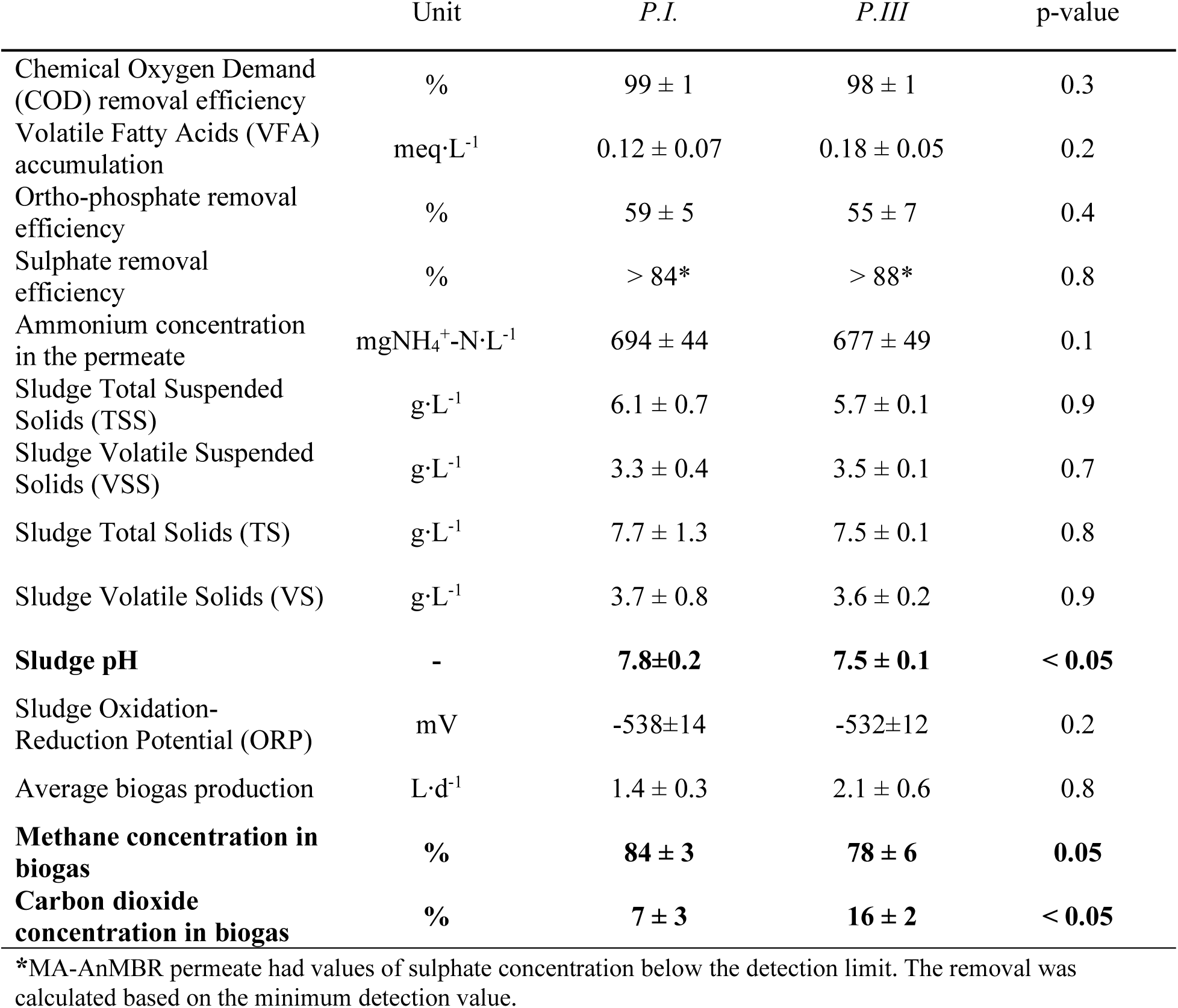
Summary of the MA-AnMBR performance during the operational pphases P.I., where the feed of the MA-AnMBR was synthetic concentrated wastewater (days 1 to 90), and P.III (days 130 to 250), when antibiotics were added to the synthetic feed. Values correspond to averages and standard deviations of samples (in triplicates). The stepwise addition of antibiotics SMX and TMP was done between days 90 and 150, reaching a final feed concentration of each antibiotic equivalent to 150 µg·L^-1^. Values shown in bold correspond to those which had statistically important changes between the different reactor phases, based on a single-factor ANOVA.

|  | Unit | P.I. | P.III | p-value |
| --- | --- | --- | --- | --- |
| Chemical Oxygen Demand (COD) removal efficiency | % | 99 ± 1 | 98 ± 1 | 0.3 |
| Volatile Fatty Acids (VFA) accumulation | meq·L<sup>-1</sup> | 0.12 ± 0.07 | 0.18 ± 0.05 | 0.2 |
| Ortho-phosphate removal efficiency | % | 59 ± 5 | 55 ± 7 | 0.4 |
| Sulphate removal efficiency | % | > 84* | > 88* | 0.8 |
| Ammonium concentration in the permeate | mgNH<sub>4</sub><sup>+</sup>·N·L<sup>-1</sup> | 694 ± 44 | 677 ± 49 | 0.1 |
| Sludge Total Suspended Solids (TSS) | g·L<sup>-1</sup> | 6.1 ± 0.7 | 5.7 ± 0.1 | 0.9 |
| Sludge Volatile Suspended Solids (VSS) | g·L<sup>-1</sup> | 3.3 ± 0.4 | 3.5 ± 0.1 | 0.7 |
| Sludge Total Solids (TS) | g·L<sup>-1</sup> | 7.7 ± 1.3 | 7.5 ± 0.1 | 0.8 |
| Sludge Volatile Solids (VS) | g·L<sup>-1</sup> | 3.7 ± 0.8 | 3.6 ± 0.2 | 0.9 |
| **Sludge pH** | - | **7.8±0.2** | **7.5 ± 0.1** | **< 0.05** |
| Sludge Oxidation-Reduction Potential (ORP) | mV | -538±14 | -532±12 | 0.2 |
| Average biogas production | L·d<sup>-1</sup> | 1.4 ± 0.3 | 2.1 ± 0.6 | 0.8 |
| **Methane concentration in biogas** | % | **84 ± 3** | **78 ± 6** | **0.05** |
| **Carbon dioxide concentration in biogas** | % | **7 ± 3** | **16 ± 2** | **< 0.05** |

\*MA-AnMBR permeate had values of sulphate concentration below the detection limit. The removal was calculated based on the minimum detection value.

The reactor pH, system average biogas production, and biogas methane concentration had statistical differences between the values obtained at the studied phases. The sludge pH decreased from 7.8 to 7.5 after the addition of antibiotics. Furthermore, while biogas production remained unchanged, around 1,400 mL·d^-1^, the biogas quality based on the methane content decreased, from 84 to 78 %. Whilst the CH_4_ concentration decreased, the carbon dioxide biogas concentration doubled, from 7 to 16%. No significant differences in sludge concentration, either suspended or total, was observed. Similarly, the nutrient content in the reactor permeate (as NH_4_^+^ and PO_4_^3-^) remained unchanged after the addition of 150 µg·L^-1^ of SMX and TMP.

Antibiotics SMX and TMP removal was assessed during phase *P.II*, where the antibiotics were introduced stepwise until a concentration of 150 µg·L^-1^ each (during a period of 40 days in total), and during phase *P.III*, for 120 days. During *P.II,* TMP concentration in the MA-AnMBR permeate remained below 10 µg·L^-1^, and SMX values were below 20 µg·L^-1^. Once stable conditions were achieved at *P.III*, the SMX and TMP removal of the MA-AnMBR was 86 ± 5% and 97 ± 1% respectively. Antibiotics concentrations adsorbed in the MA-AnMBR sludge were similar to the ones found in the permeate, 9 ± 4 µg·L^-1^ and 14 ± 6 µg·L^-1^ of TMP and SMX respectively.

### 3.2. Antibiotics adsorption batch tests

Batch tests with MA-AnMBR sludge, taken from the reactor during *P.I*, were conducted to assess TMP and SMX adsorptions at 10 °C, for concentrations of 10, 50, and 150 µg·L^-1^.

For all studied concentrations, the adsorbed TMP fraction was around 82% after six hours, as shown in Figure 3. TMP had an elevated level of adsorption in the first five minutes of testing. TMP concentrations in the liquid after 5 minutes were below 30% of the initial concentrations applying 10 and 50 µg·L^-1^, and 50% for the initial concentration of 150 µg·L^-1^. A single-factor ANOVA statistical analysis was conducted to assess the adsorption differences between the different initial TMP concentrations (10, 50, and 150 µg·L^-1^). No statistical differences were found after 6 hours of testing (p-value of 0.2).

**Figure 3.**
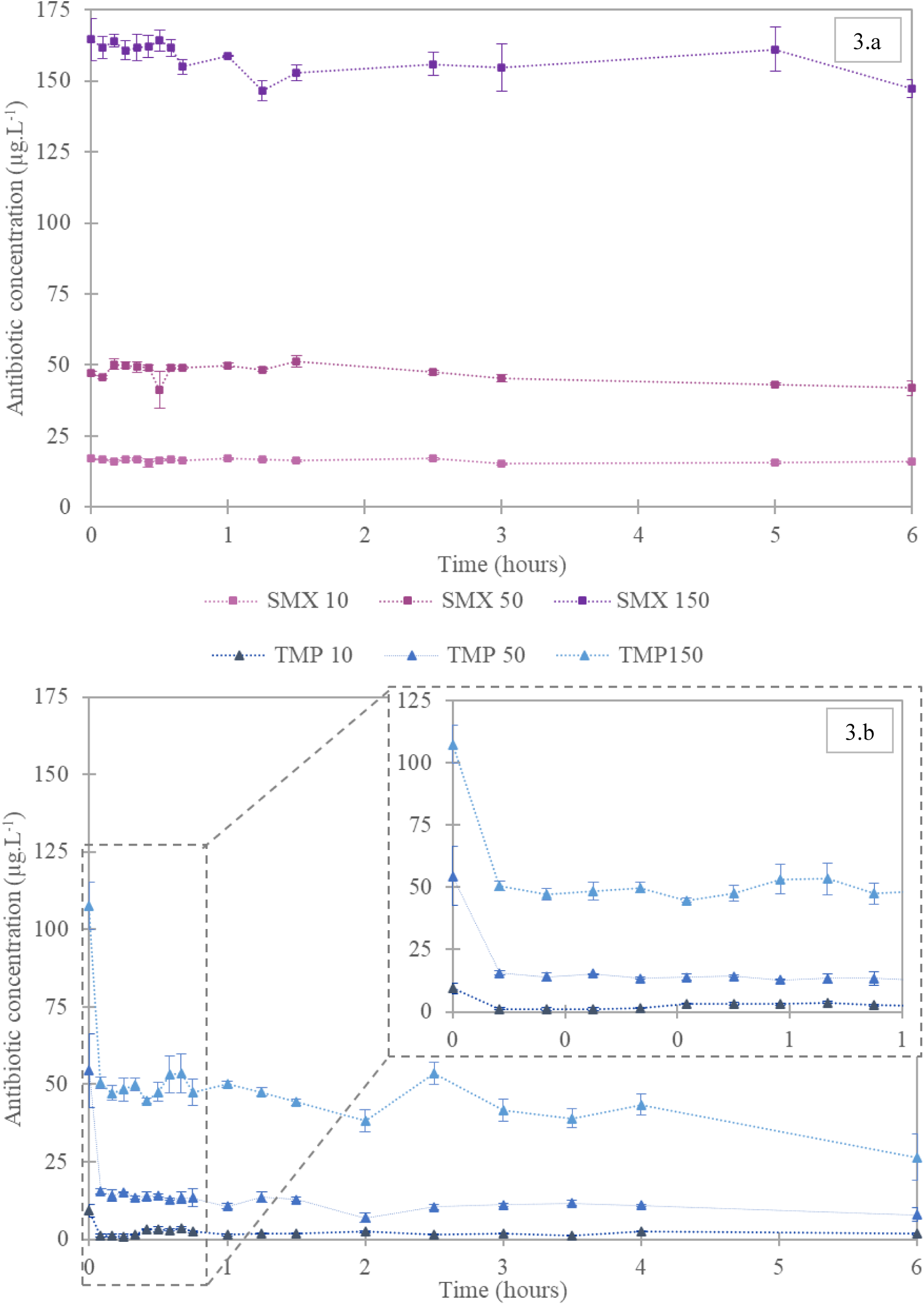
Adsorption batch tests of antibiotics SMX (Figure 3.a) and TMP (Figure 3.b) at 10°C, with MA-AnMBR sludge. The sludge total solids concentration was 4.1 and 3.9 g·L^-1^ for SMX and TMP samples, respectively. Additionally, a total volume of around 100 mL of sludge was used for each antibiotic concentration, and experiments were performed in triplicates.

Adsorption of SMX onto the MA-AnMBR sludge was minimal, with values of 11% after six hours of testing (Figure 3). No statistical differences were found (p-value of 0.4) between the observed adsorption at the different SMX concentrations.

### 3.3. Antibiotic-resistant bacteria

The levels of ARB in the MA-AnMBR bulk and permeate were measured by heterotrophic plate count in three out of the four study phases: *P.II*, *P.III*, and *P.IV*, while the total bacterial concentration was additionally measured during phase *P.I* (using the methods described in 2.4.2). Results are shown in Figure 4. Total bacteria removal in the MA-AnMBR system was in the order of 3 log in all studied phases, 99.9% (difference between the bacteria count in the reactor bulk and the UF permeate). Removal of ARB varied and depended on the experimental phase. No bacteria resistant to SMX or TMP were found in the permeate after 21 days of TMP supply (day 111; influent TMP concentration was 150 µg·L^-1^ and SMX was not yet added).

**Figure 4.**
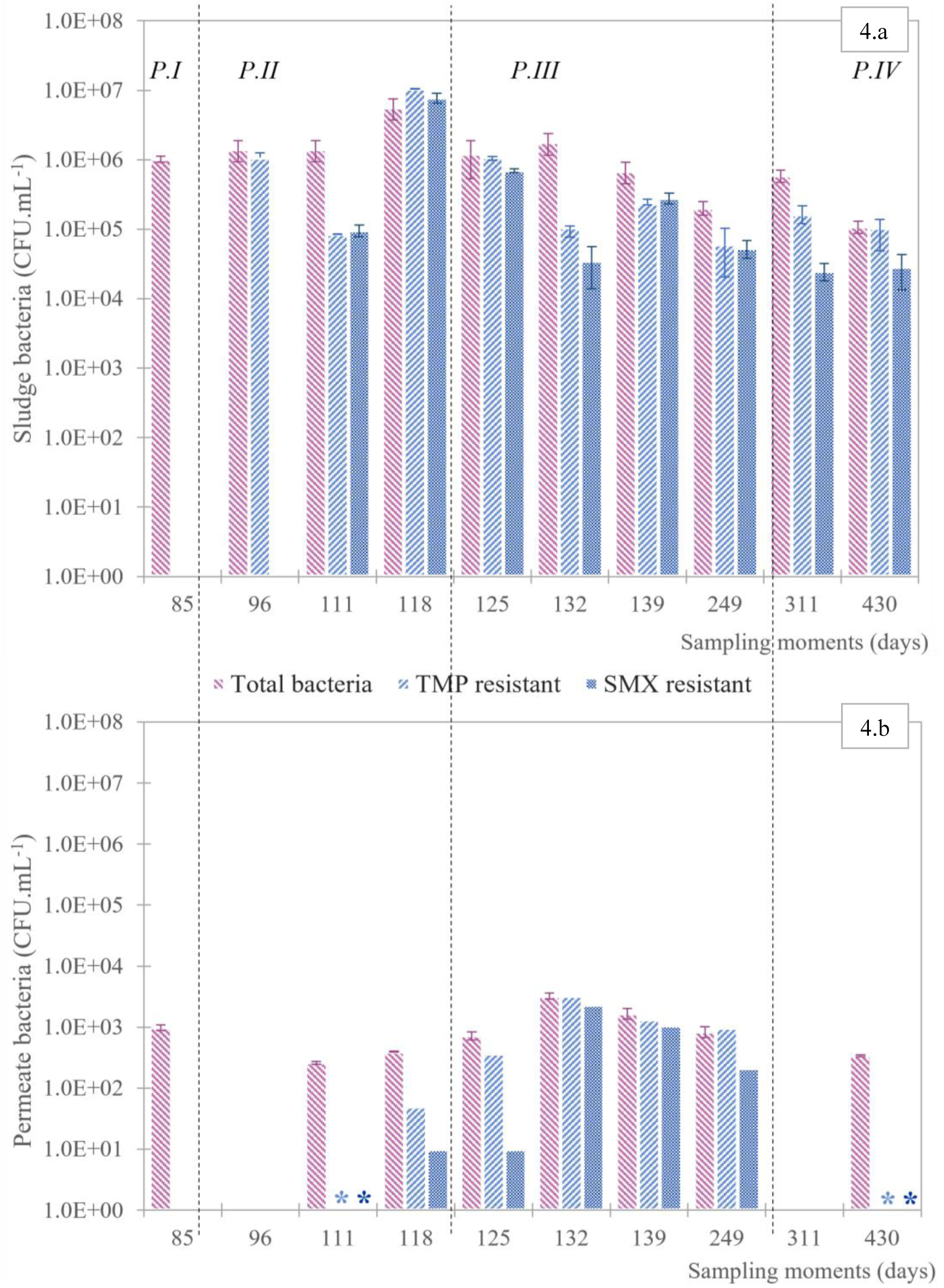
Antibiotic-resistant and total bacteria in the MA-AnMBR. Fig. 4.a shows the MA-AnMBR reactor bulk bacteria. Fig. 4.b shows the MA-AnMBR UF permeate bacteria. The reactor phases are visualized with dotted vertical lines. P. I corresponds to the phase before antibiotics were added to the MA-AnMBR synthetic feed; P.II is referred to as the phase when the antibiotics SMX and TMP were added in steps, in the concentration of 10, 50 and 150 µg·L^-1^. Each step lasted around 3 HRT (7.5 days). P.III phase lasted 120 days in which SMX and TMP were present in the MA-AnMBR feed with a concentration equivalent to 150 µg·L^-1^each. Finally, P.IV refers to the phase where the influent of the MA-AnMBR had no antibiotics. Days with * refer to measured samples with values below the detection limit. Blank days indicate no measurements.

During *P.IV*, the MA-AnMBR influent was supplied again with a feed without antibiotics. No bacteria resistant to SMX nor TMP were detected anymore from the MA-AnMBR permeate. SMX-resistant bacteria followed a similar trend to TMP-resistant bacteria. SMX-resistant ones were removed by 5 log during *P.II* and by 2 log during *P.III*. Likewise, TMP-resistant bacteria were removed by 4 log during *P.II* and by 2 log during *P.III*. Furthermore, during *P.III*), the concentration of total bacteria in the permeate was similar to the ARB ones.

### 3.4. Antibiotic resistance genes

The ARGs *sul1*, *sul2* and *dfrA1*, and the MGE *intI1* were measured during the four reactor phases from the MA-AnMBR sludge. All genes were already found in the sludge sample that was taken before the addition of the antibiotics, on day 90 of operation. During *P.II*, all four genes showed an increase in their relative abundance (per 16S rRNA gene), as shown in Figure 5. A peak in all four gene relative abundances was observed on day 112, which corresponded to the start of the addition of 10 µg·L^-1^ SMX.

**Figure 5.**
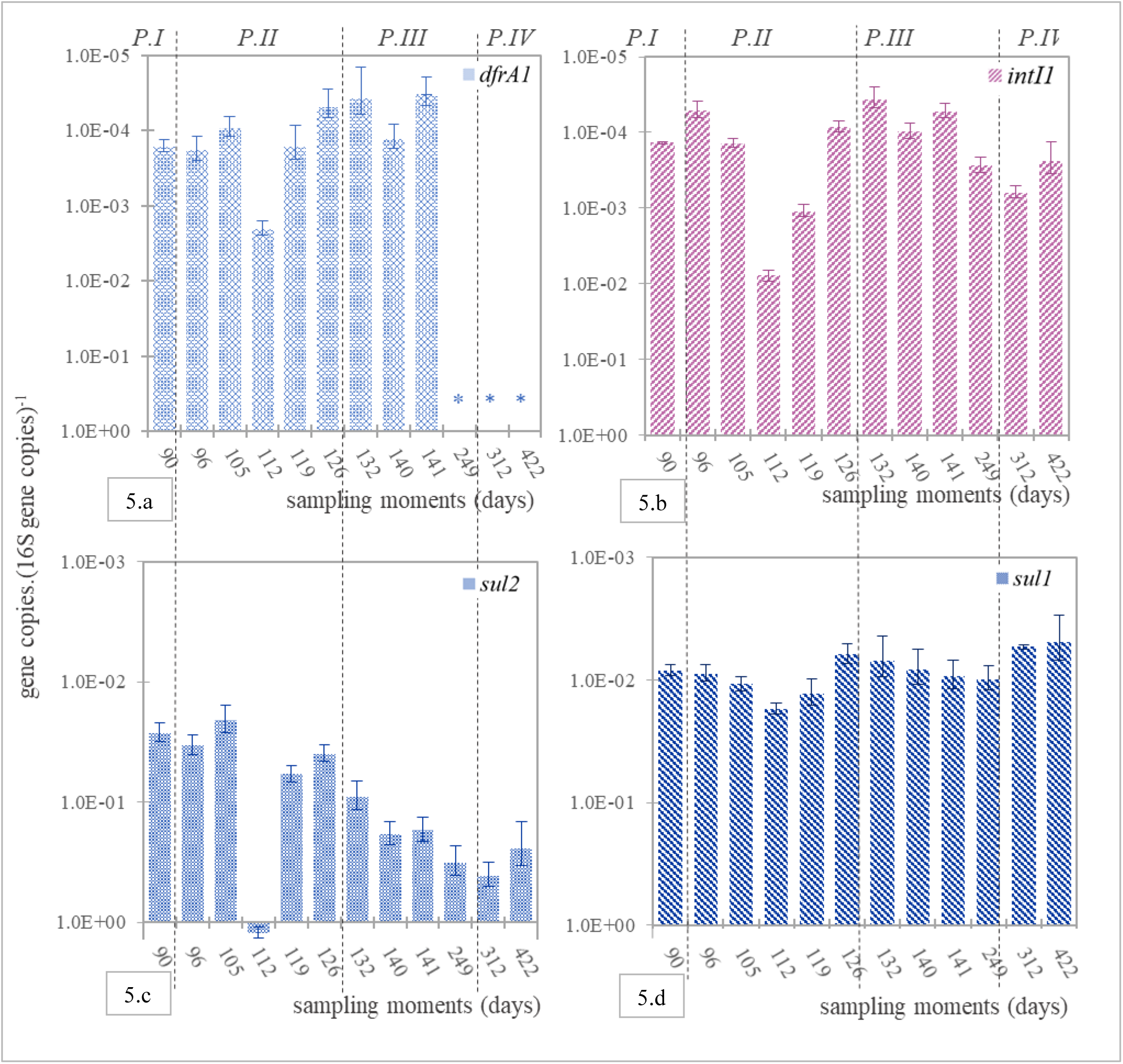
Antibiotic-resistant gene copies per 16S gene copies of the MA-AnMBR sludge. Fig. 5.a refers dfrA1 gene concentration. Fig. 5.b refers to intI1 concentration. Fig. 5.c. refers to sul2 concentration, and, Fig. 5.d. refers sul1 gene concentration. The reactor phases are visualized with dotted vertical lines. P. I corresponds to the phase before antibiotics were added to the MA-AnMBR synthetic feed; P.II is referred to as the phase when the antibiotics SMX and TMP were added in steps, in the concentration of 10, 50 and 150 µg·L^-1^. Each step lasted around 3 HRT (7.5 days). P.III phase lasted 120 days in which SMX and TMP were present in the MA-AnMBR feed with a concentration equivalent to 150 µg·L^-1^each. Finally, P.IV refers to the phase where the influent of the Ma-AnMBR had no antibiotics. Values below the detection limit for dfrA1 (5.2E3 dfrA1 gene copies) are presented with *, and blank days indicate no measurements.

Relative abundances of the *sul2* gene were highest among the analyzed genes, with an average difference corresponding to two orders of magnitude. Aside from the peak on day 112, *sul2* increased during *P.II* and *P.III* to a value of 3.2 ×10^-1^ gene copies·(16S gene copies)^-1^ at the end of *P.III* (day 249). During *P.IV*, the dosing of antibiotics in the influent was stopped, however, after 62 days in *P.IV* (day 312), *sul2* reached even 4.1 ×10^-1^ gene copies· (16S gene copies)^-1^. This concentration decreased to only 2.4×10^-1^ gene copies· (16S gene copies)^-1^ on the last day of reactor operation (day 422). A reduction in gene copies in phase *P.IV* was also observed for *dfrA1* and *sul1*, indicating a loss of antibiotic-resistance genes when antibiotics dosage to the MA-AnMBR was stopped. Concentrations of *sul1* during *P.IV* were even found to be below the values measured before the start of the antibiotic’s addition: 5.0 x10^-3^ and 8.4 x10^-3^ gene copies.(16S gene copies)^-1,^ respectively. Finally, gene copies were below the detection limit of 5.2 x10^-3^ for *dfrA1* at the end of *P.III* (day 249 days).

Gene copies from the UF permeate of the MA-AnMBR were measured during the first three reactor phases. UF permeate *dfrA1* gene copies were below the detection limit in all samples. For the rest of the studied genes, a difference of one to four orders of magnitude was found between the total abundance of gene copies (per mL of sample) in the bulk and the ones in the MA-AnMBR UF permeate, as shown for *sul1* in Figure 6. Apparently, the UF membrane of the MA-AnMBR retained the majority of the microorganisms that contained the studied genes, reducing 99.9% of the studied genes. Nevertheless, when the relative abundance of gene copies are considered, the concentration of copies in the UF permeate and the reactor bulk tends to be in the same order of magnitude, as shown for the *sul1* gene in Figure 6. Permeate and reactor bulk concentrations of gene *sul2* and MGE *intI1* can be found in **Annex C.**

**Figure 6.**
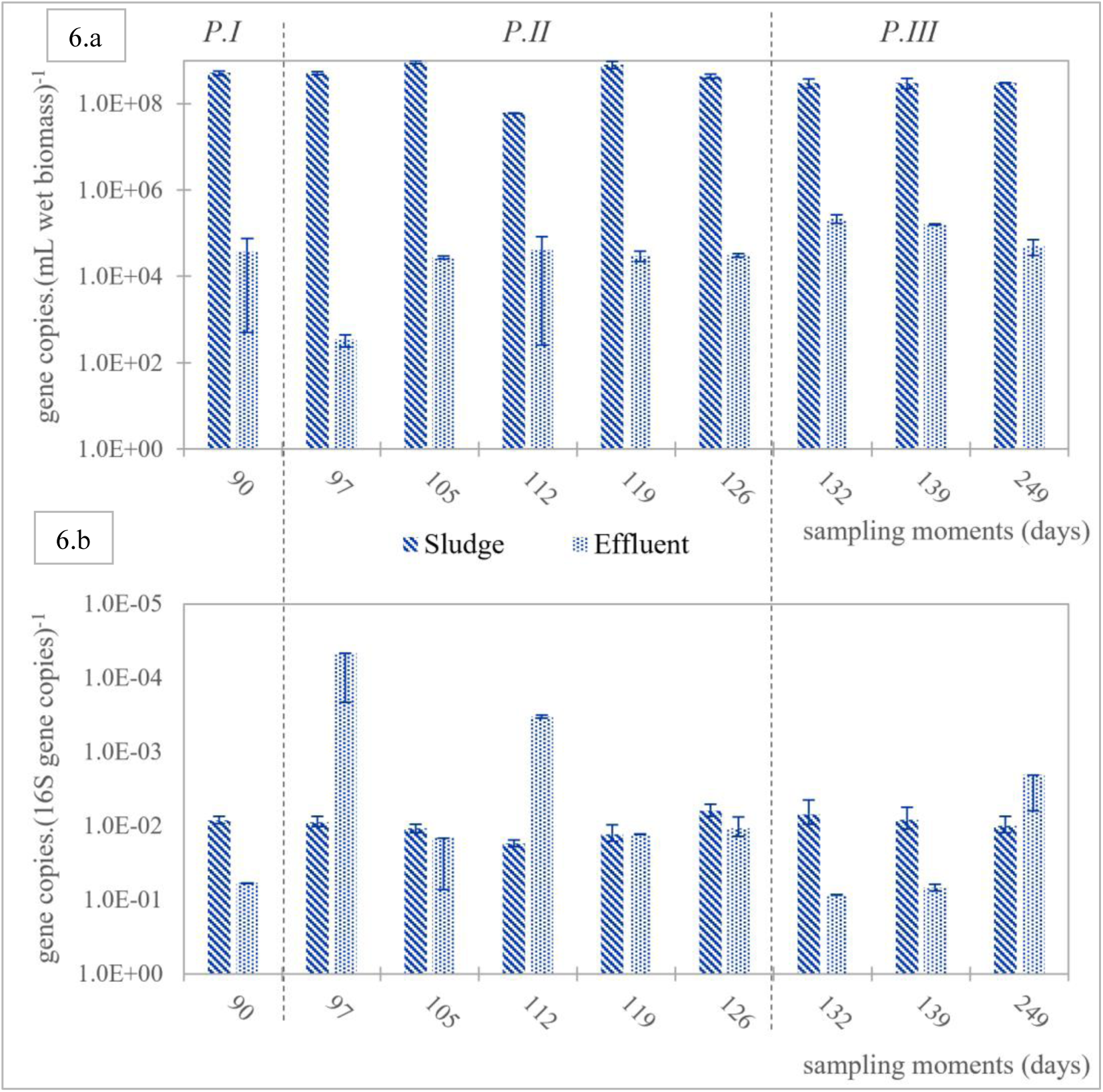
The concentration of gene sul1 in the MA-AnMBR sludge and permeate. Fig. 6.a shows to the total gene copies per mL of wet biomass, while the values in Fig.6.b show values of sul1 gene standardized per 16S gene copies. The reactor phases are visualized with dotted vertical lines. P. I corresponds to the phase before antibiotics were added to the MA-AnMBR synthetic feed; P.II is referred to as the phase when the antibiotics SMX and TMP were added in steps, in the concentration of 10, 50 and 150 µg·L^-1^. Each step lasted around 3 HRT (7.5 days). P.III phase lasted 120 days in which SMX and TMP were present in the MA-AnMBR feed with a concentration equivalent to 150 µg·L^-1^each. Finally, P.IV refers to the phase where the influent of the Ma-AnMBR had no antibiotics.

## 4. Discussion

### 4.1. Removal and consequences of TMP and SMX additions on the MA-AnMBR

Despite the addition of air in the MA-AnMBR, TMP removal was high, i.e., 97 ± 1%. A similar TMP removal of around 94% was obtained in a lab-scale AnMBR treating synthetic sewage [27]. Furthermore, Feng, et al. [49] have found that TMP removal on anaerobic digestion of pig manure was above 99% after 10 days of digestion. In contrast, a TMP removal of only 26.4% has been measured from conventional activated sludge processes, that are mostly operated with aerobic conditions or in alternance of redox conditions [50]. Previously, the changes in AnMBR performance with and without micro-aeration had been assessed: it was concluded that the given micro-aeration (at to an oxygen over influent COD load of 1.0%) had a negligible impact on the reactor performance (Piaggio et al, submitted). Thus, based on the efficiency measured in the MA-AnMBR, it can be inferred that the micro-aeration of the system has a negligible effect on TMP removal, and the MA-AnMBR is efficient in the removal of this antibiotic.

The log K_ow_ value of TMP and its positive charge at circumneutral pH results in quick adsorption onto the negatively charged biomass [51]. Adsorption of TMP at 10 °C in batch tests showed a removal of TMP around 75% after 6 hours of testing, as shown in Figure 3. In the continuous-flow MA-AnMBR, removal of TMP from the permeate reached 97%. From the results obtained in the adsorption tests, the sludge adsorption capacity for TMP was around 200 μg·gTSS. The MA-AnMBR has a TSS content of around 6g·L^-1^ and total volume of 6.5 L, resulting in a total biomass suspended biomass of 39 g. Considering the above-mentioned adsorption capacity, the MA-AnMBR is expected to be TMP saturated after 21 days of operation. Hereafter, if no biodegradation of TMP occurred, TMP should be accumulated, and its concentration should increase. Since the removal of TMP in the MA-AnMBR is 97%, it can be concluded that TMP was degraded in the MA-AnMBR. Antibiotic concentration in the MA-AnMBR sludge was measured during *P.II* and *P.III*. The residual TMP concentration increased from *P.II* to *P.III* until it reached a plateau, with a concentration of 9 ± 4 µg·L^-1^. Thus, TMP is quickly adsorbed onto the sludge and highly likely subsequently digested. Apparently, the applied SRT of 27 days allowed enough time for the anaerobic degradation of TMP. Alvarino, et al. [17], Feng, et al. [49].

The removal of SMX in the MA-AnMBR was 86 ± 5%. This value was in the same range, between 70 to 90%, of reported SMX removal in fully anaerobic lab-scale AnMBR units treating domestic wastewater [45, 52, 53]. SMX removal in an aerobic activated sludge process has been reported to be much lower than in the MA-AnMBR, i.e., 39.1% [50]. ORP measurements showed that anaerobic conditions were kept in the MA-AnMBR even under micro-aeration, maintaining an ORP of -530 mV. Whilst a high removal of SMX was obtained in the MA-AnMBR, batch adsorption tests at 10 °C showed SMX removal below 15% (Figure 3). The low adsorption of SMX onto the sludge is likely due to the negative charge of SMX at circumneutral pH, disfavoring its attraction to negatively charged biomass. The residual SMX concentration in the MA-AnMBR sludge was only 14 ± 6 µg·L^-1^ during *P.III*.

Most likely SMX was also degraded anaerobically in the MA-AnMBR, considering the measured removal efficiencies. With the measured low adsorption, the degradation rate of SMX is determined by the hydraulic loading rate and the HRT of 2.6 days. The most important transformation reactions of the isoxazole ring of SMX and routes to co-metabolize SMX are hydroxylation, hydrogenation, acetylation, desulfurization and reductive cleavage, among others [54, 55]. Some of the degradation pathways of SMX, like hydroxylation and desulfurization as part of acetylation, take less than one day, ensuring SMX degradation under short HRT [55]. According to Dermer and Fuchs [56], dehydrogenases enzymes play a key role in the hydroxylation of SMX under anaerobic conditions, and hydroxylation can be considered one of the main SMX degradation pathways. Therefore, while TMP is rapidly adsorbed into the MA-AnMBR sludge, and then the long SRT of the systems provides enough time for its degradation, SMX is barely adsorbed into the sludge, but the applied HRT of 2.6 days of the laboratory-scale MA-AnMBR provides enough time for its rapid degradation. Thus, applied HRT of 2.6 days could be considered as a design criterion for the removal of TMP.

While biogas quality changes from *P.I* to *P.III* were statistically significant, no change in biogas quantity was observed. Cetecioglu, et al. [57] have shown that only concentrations above 45 mg·L^-1^ of SMX were lethal to the microbial community and hence, inhibit biogas production. Zarei-Baygi, et al. [45] have concluded that after the addition of 250 µg·L^-1^ of SMX to a laboratory-scale AnMBR fed with synthetic wastewater, there was no significant difference in the abundance of methanogens, and the microbial community of biomass was stable throughout the study. Furthermore, Tang, et al. [55] concluded that addition of up to 2 mg·L^-1^ of SMX was beneficial for methane production, since the addition of SMX impacted the acetogenesis and therefore, shortened the time required to reach the maximum methane production.

### 4.2. ARB and ARG

The results of this long-term experiment indicate a potential development of antibiotic resistance in the bacterial population when antibiotics are present in the wastewaters. On the Ma-AnMBR permeate, no ARB bacteria were present in day 111 and 430. The latter sampling moment was performed when the MA-AnMBR was fed with synthetic feed an no antibiotics, while the samples taken on day 111 corresponded to the moment the Ma-AnMBR was fed with the synthetic feed and 150 µg·L^-1^ of TMP. ARB measurements executed after day 111 showed an increase in both TMP and SMX relative concentrations (to the total heterotrophic bacteria count), as shown in Figure 4. The highest concentration of TMP and SMX ARB were in the UF permeate were obtained on *P.III*, having a relative abundance of 57 and 70% respectively. Furthermore, TMP ARB in the Ma-AnMBR bulk increased from 4% of the total heterotrophic bacteria count at *P.II*, to 9% at *P.III* and finally 20% at *P.IV*. On the other hand, no significant difference was found (p value of 0.8) in the relative abundance of the TMP and SMX ARB in the reactor bulk during the studied periods. Thus, addition of the antibiotics affected the TMP ARB but not the SMX ARB in the MA-AnMBR mixed liquor.

Both SMX and TMP-resistant bacteria were measured in the MA-AnMBR permeate during phases *P.I* to *P.III,* but no resistant bacteria were found 180 days after the antibiotic’s dosage to the MA-AnMBR was stopped (day 430, *P.IV*). Whilst these results might indicate the loss of resistance, antibiotic-resistant genes for SMX were found in abundance in the MA-AnMBR permeate, indicating the contrary. The antibiotic-resistant gene chosen for TMP, *dfrA1*, was below the detection limit in all phases of permeate samples. Thus, the resistance towards this antibiotic in the MA-AnMBR should not be measured based on the selected ARG, but on the ARB count.

The MA-AnMBR was equipped with a UF membrane with a nominal pore size of 30 nm, which was apparently too big for complete removal of total bacteria or ARB, under all the studied phases. Lousada-Ferreira, et al. [58] have found that particles 100 times bigger than the nominal pore size were found in several membranes’ permeates, even when membrane integrity was not compromised. The authors concluded that the nominal membrane pore size given by the manufacturer is rather an indication of the average membrane pore size but might be very different from the maximum values. Most municipal wastewater bacteria sizes vary between 1 to 100 µm, for example, *E.coli* has an average size of 1-2 µm [59]. Therefore, it is possible that some bacteria will pass the UF membrane of the MA-AnMBR. Furthermore, the permeate of the laboratory-scale MA-AnMBR is rich in nutrients, such as nitrogen and phosphorus, and has an optimal temperature for bacterial re-growth. Therefore, the determined values around 10^3^ CFU·mL^-1^ were expected for the laboratory-scale MA-AnMBR permeate, representing a 3 log removal of total bacteria and ARB.

All measured ARGs and *intI1* relative abundances in the MA-AnMBR sludge and UF permeate increased during phase *P.II*, but only *sul1* and *sul2* in the sludge kept increasing during the phase *P.III*, as shown in Figure 5. This results are aligned with the results obtained by Guo, et al. [60], that found an increase in *intI1* mobile gene element, *sul1* and *sul2* due to an addition of oxygen equivalent to 1% of influent COD, in a sequencing batch reactor fed with blackwater. The increase in *sul1* and *sul2* genes due to the addition of SMX, has also been observed by Zarei-Baygi, et al. [24] and Blahna, et al. [61]. Whilst SMX resistance genes increased during *P.II* and *P.III,* no statistical differences were found in the relative concentration of SMX-resistant bacteria and total bacteria between these phases (p-value of 0.4). Furthermore, the relative abundance of ARG in the permeate was similar to the one obtained in the sludge. Thus, while the UF membrane can retain 99.9% of the measured bacteria, the bacteria present in the permeate has a similar abundance of ARGs.

Pearson correlation tests were conducted between the measured levels of SMX-resistant bacteria and TMP-resistant bacteria, and the relative abundances of the different genes, for samples taken from the reactor bulk and the permeate of the MA-AnMBR. A strong linear correlation was assumed when the absolute value of the Pearson coefficient (ρ) was above 0.7. A strong linear positive correlation was observed between TMP- and SMX-resistant bacteria in the reactor bulk and UF permeate, with ρ values of 0.83 and 0.97 respectively (**Annex D**). Thus, SMX-resistant bacteria increased simultaneously with TMP-resistant bacteria, which might be attributed to the fact that both antibiotics were more or less simultaneously added to the MA-AnMBR, with only 20 days difference between the start of dosing TMP and SMX.

The gene *dfrA1* was also positively correlated to all studied genes on the sludge samples of the MA-AnMBR. When a concentration of 10 µg·L^-1^ of SMX started to be added to the reactor on day 112, all studied genes reached their maximum concentration (Figure 5). Thus, the addition of SMX is not only linked to the concentration of the genes associated with this antibiotic (*sul1* and *sul2*). A change in *dfrA1* gene concentration could be linked to both SMX and/or TMP addition, and therefore, TMP resistance should not be considered only based on changes in *dfrA1* concentration. Results showed that plate counts of resistant bacteria is a better estimation for measuring resistance towards TMP than the relative abundance of the *dfrA1* gene.

Finally, all measured relative abundances of the *sul1*, *sul2*, *dfrA1*, and *intI1* genes decreased during phase *P.*IV, i.e., when the MA-AnMBR was fed without antibiotics. Thus, biological digestion in the MA-AnMBR is favourable for the reduction of ARG levels in the sludge. Several authors have stated that biological treatment and membrane systems are efficient for the removal of ARGs, especially *sul1* and *sul2* [24, 62, 63]. Nevertheless, considering the relative abundance of genes (vs. 16S rRNA gene copy number), the UF membrane system of the MA-AnMBR might not retain all microorganisms, and the ones passing the membrane still have ARGs. Thus, while the biological treatment achieved an increased removal of ARGs, it is not efficient in retaining all microorganisms, and similar ARGs abundance to the one in the reactor bulk can be found in the UF permeate. Thus, for the removal of ARB and microorganisms containing the studied ARGs, a following treatment step should be considered.

## 5. Conclusions

The fate of the SMX and TMP antibiotics was studied on a laboratory scale micro-aerated AnMBR fed with synthetic concentrated domestic wastewater. Antibiotic resistance spreading was assessed by measuring the levels of ARB, ARGs *sul1, sul2, dfrA1*, and MGE *intI1* in the sludge and permeate. The effect of the additions of the antibiotics on the performance of the MA-AnMBR, their removal, and their relation to resistance induction were assessed for 430 days. The main conclusions of the research are the following:

- The addition of 150 µg·L^-1^ of SMX and TMP into the MA-AnMBR feed had negligible effects on the system performance. The sludge pH decreased from 7.8 to 7.5, which simultaneously entailed an increase of CO_2_ concentration in the biogas (from 7 to 16%) and a decrease in CH_4_ partial pressure. These changes were statistically significant (p-value <0.05).
- A high removal of SMX and TMP was achieved in the laboratory-scale MA-AnMBR. SMX was poorly adsorbed into the sludge but rapidly degraded, reaching a total removal of 86%, measured in the MA-AnMBR permeate. In contrast, TMP was rapidly adsorbed onto the MA-AnMBR sludge and the long SRT of the system promoted its degradation, achieving a total TMP removal of 97%. Thus, micro-aeration of the membrane system had no negative effect on the removal of the antibiotics.
- ARB and ARGs were found in both the MA-AnMBR sludge and permeate. While the system was able to reduce the ARB concentration by 3 log, the ARGs abundance (relative to the 16S-rRNA gene) was similar in the sludge and the ultrafiltration permeate. The addition of SMX and TMP led to an increase in the relative abundance of all ARGs and *intI1* in the MA-AnMBR sludge.
- The relative abundance of the ARG *dfrA1* in the MA-AnMBR mixed liquor had a strong linear correlation with *sul1*, *sul2*, and the MGE *intI1.* However, changes in relative abundance of the genes were not linked ARB. Thus, the gain of resistance to TMP or SMX is better assessed by the heterotrophic plate count of ARB than by molecular detection of the genes.
- The relative abundance of the MGE *intI1* in the sludge was positively and linearly correlated with all measured genes (ρ values above 0.82), reflecting the overall antibiotic pressure. It also suggests that microorganisms potentially mobilized their ARG pool to resist.

## Acronyms and symbols

AnMBR: Anaerobic membrane bioreactor
ARB: Antibiotic-resistant bacteria
ARG: Antibiotic-resistant gene
COD: Chemical oxygen demand
GC: Gas chromatography
HGT: Horizontal gene transfer
HRT: Hydraulic retention time
KOW: Octanol-water partition coefficient
MA-AnMBR: Micro-aerated anaerobic membrane bioreactor
MBR: Membrane bioreactor
MGE: Mobile genetic elements
ORP: Oxidation-reduction potential
qPCR: Quantitative polymerase chain reaction
SMX: Sulfamethoxazole
SRT: Solids retention time
TMP: Trimethoprim
TS: Total solids
TSS: Total suspended solids
UF: Ultrafiltration
VFA: Volatile fatty acids
VS: Volatile solids
VSS: Volatile suspended solids
WWTP: Wastewater treatment plant

## Acknowledgements

This research was funded by NWO in the Netherlands (Project number 15424-2) and the Department of Biotechnology (DBT) in India (BT/IN/Indo-Dutch/19/TRS/2016), as part of the LOTUS-HR program (https://lotushr.org). Our gratitude goes to the lab technicians at the TU Delft Water Lab. Furthermore, we are thankful for the work done by Srilekha Mittapalli and Pravin Khande on the MA-AnMBR during their time at the TU Delft. The authors are grateful for Julia Gebert’s contributions and supervision.

