## Supplementary Material for "The fate of sulfamethoxazole and trimethoprim in a micro-aerated anaerobic membrane bioreactor: implications for antibiotic resistance spreading"

### A. Lab-scale MA-AnMBR reactor conditions

***Table A.1.*** *Micro-aerated AnMBR feed recipe.*

| Feed composition | Unit | Value | Micronutrients Solution | Unit | Value |
|---|---|---|---|---|---|
| Urea | $g.L^{-1}$ | 1.0 | $FeCl_3.6H_2O$ | $mg.L^{-1}$ | 1000.0 |
| Ammonium chloride | $g.L^{-1}$ | 0.8 | $CoCl_2.6H_2O$ | $mg.L^{-1}$ | 1000.0 |
| Sodium acetate trihydrate | $g.L^{-1}$ | 2.6 | $MnCl_2.4H_2O$ | $mg.L^{-1}$ | 250.0 |
| Ovalbumin | $g.L^{-1}$ | 0.2 | $CuCl_2.2H_2O$ | $mg.L^{-1}$ | 15.0 |
| Magnesium sulphate heptahydrate | $g.L^{-1}$ | 0.1 | $ZnCl_2$ | $mg.L^{-1}$ | 25.0 |
| Potassium phosphate monobasic | $g.L^{-1}$ | 0.2 | $H_3BO_3$ | $mg.L^{-1}$ | 25.0 |
| Calcium chloride dihydrate | $g.L^{-1}$ | 0.1 | $(NH_4)_6Mo_7O_{24}.4H_2O$ | $mg.L^{-1}$ | 45.0 |
| Cellulose | $g.L^{-1}$ | 1.5 | $Na_2SeO_3.H_2O$ | $mg.L^{-1}$ | 50.0 |
| Milk powder | $g.L^{-1}$ | 0.6 | $NiCl_2.6H_2O$ | $mg.L^{-1}$ | 25.0 |
| Yeast extract | $g.L^{-1}$ | 0.5 | EDTA | $mg.L^{-1}$ | 500.0 |
| Sunflower oil | $drops.L^{-1}$ | 2.0 | HCI 36% | $mg.L^{-1}$ | 0.5 |
| Humic and Fulvic acid | $drops.L^{-1}$ | 2.0 | Resazurin sodium salt | $mg.L^{-1}$ | 250.0 |
| Micronutrients solution | $g.L^{-1}$ | 10.6 | Yeast extract | $mg.L^{-1}$ | 1000.0 |

***Table A.2.*** *. Micro-Aerated AnMBR feed composition*

| | Unit | Value |
|---|---|---|
| Chemical Oxygen Demand (COD) | $mg.L^{-1}$ | 5200 ± 600 |
| Ammonium ($NH_4^+$) | $mgN.L^{-1}$ | 249± 54 |
| Nitrate ($NO_3^-$) | $mgN.L^{-1}$ | 1.3 ± 0.2 |
| Phosphate ($PO_4^{3-}$) | $mgP.L^{-1}$ | 60 ± 9 |
| Sulphate ($SO_4^{2-}$) | $mgS.L^{-1}$ | 235 ± 46 |
| Total Suspended Solids (TSS) | $mg.L^{-1}$ | 3073 ± 451 |
| Volatile Suspended Solids (VSS) | $mg.L^{-1}$ | 2938 ± 436 |

### B. Mix solution and reaction conditions for qPCR

All ARGs and *intl-1* qPCR reactions were conducted using a master mix per sample. The master mix consisted of a total volume of 20 µL, including IQ™ SYBR green supermix BioRad 1x, of which 0.4 µL were of each forward and reverse primer (50 µM), 10 µL of SYBR green dye, 7.6 µL of qPCR grade water, and 2 µL of the DNA template. All the reactions were performed in technical triplicates, using a qTOWER3 Real-time PCR machine (Westburg, DE). The PCR cycles and amplification conditions depended on the selected gene. **Table B-1** summarized the amplification conditions for each gene. Forward and reverse primers are summarized in **Table B-2.**

***Table B-1**. Amplification conditions per selected ARGs, MGE and 16srRNA.*

| Genes | Conditions |
|---|---|
| *sul1* | 5 minutes at 95 °C, 40 cycles of 15 seconds at 95 °C, annealing 30 seconds at 65 °C |
| *sul2* | 5 minutes at 95 °C, 40 cycles of 15 seconds at 95 °C, annealing 30 seconds at 61 °C |
| *dfrA1* | 5 minutes at 95 °C, 40 cycles of 10 seconds at 95 °C, annealing 30 seconds at 60 °C |
| *intl-1* | 5 minutes at 95 °C, 40 cycles of 15 seconds at 95 °C, annealing 30 seconds at 60 °C |
| 16s-rRNA | 5 minutes at 95 °C, 40 cycles of 15 seconds at 95 °C, annealing 30 seconds at 60 °C |

***Table B-2.** Forward and reverse primers of the selected ARGs, MGE and 16SrRNA.*

| Genes | Forward Primer 5′-3′ | Reverse Primer 5′-3′ |
|---|---|---|
| *sul1* | CGCACCGGAAACATCGCTGCAC | TGAAGTTCCGCCGCAAGGCTCG |
| *sul2* | TCCGGTGGAGGCCGGTATCTGG | CGGGAATGCCATCTGCCTTGAG |
| *dfrA1* | TTCAGGTGGTGGGGAGATATAC | TTAGAGGCGAAGTCTTGGGTAA |
| *intl-1* | GATCGGTCGAATGCGTGT | GCCTTGATGTTACCCGAGAG |
| 16s rRNA | ACTCCTACGGGAGGCAGCAG | ATTACCGCGGCTGCTGG |

Standards were added to each PCR plate to generate the standard curve. At least 6 serial dilution points (in technical duplicate) were performed to create the standard curves. An average standard curve based on the curve generated in each run was created for every gene set. Finally, the gene concentration values were then calculated from the aforementioned curve. For *sul1, sul2*, *dfrA1* and *intl-1*, gene concentration values were standardized based on the 16srRNA gene concentration of each sample (sludge or permeate).

### C. Gene concentration in the MA-AnMBR

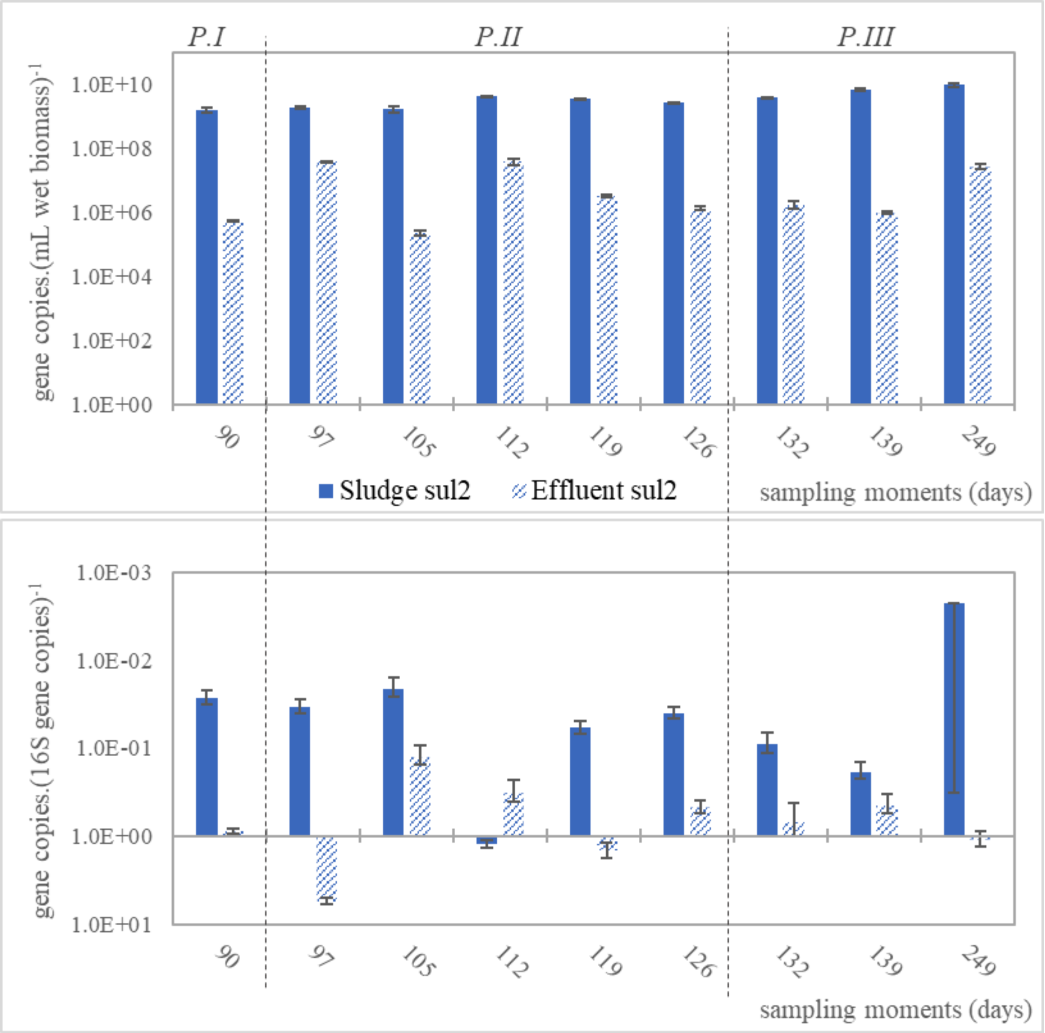


*Figure 0-1. Concentration of gene sul2 on the MA-AnMBR sludge and effluent. The figure on the top corresponds to the total gene copies per mL of wet biomass, while the values in the figure from the bottom are values of the sul2 gene standardized per 16S gene copies. The graphs show the reactor periods in which the samples were taken P.I corresponds to the period before antibiotics were added to the MA-AnMBR feed; P.II entitles the period in which the antibiotics SMX and TMP were added in steps, in the concentration of 10, 50 and 150 µg.L$^{-1}$. Each step lasted around 3 HRT (7.5 days). Finally, the P.III entitles 120 days in which SMX and TMP were present in the MA-AnMBR feed with a concentration equivalent to 150 µg.L$^{-1}$each..*

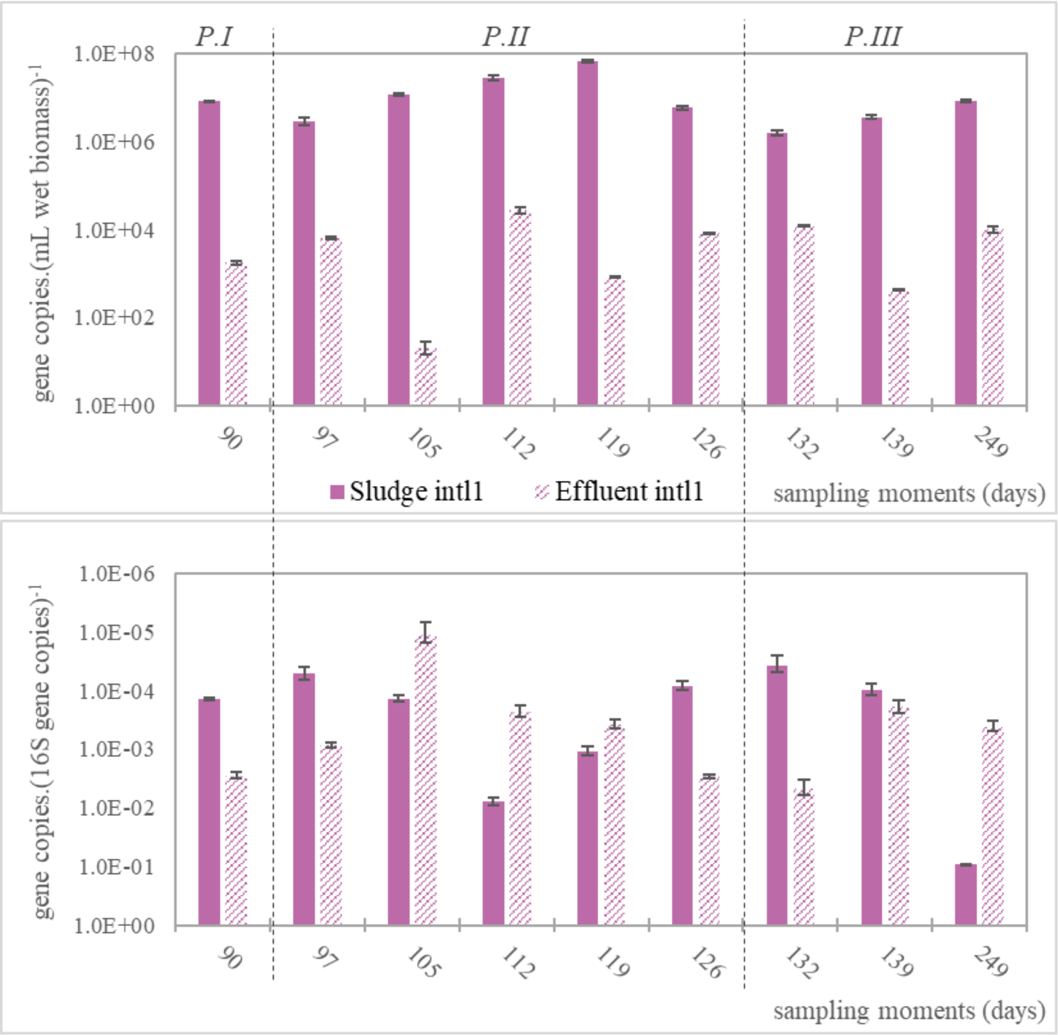


*Figure 0-2. Concentration of gene intl-1 on the MA-AnMBR sludge and effluent. The figure on the top corresponds to the total gene copies per mL of wet biomass, while the values in the figure from the bottom are values of the intl-1 gene standardized per 16S gene copies. The graphs show the reactor periods in which the samples were taken. P.I corresponds to the period before antibiotics were added to the MA-AnMBR feed; P.II entitles the period in which the antibiotics SMX and TMP were added in steps, in the concentration of 10, 50 and 150 µg.L$^{-1}$. Each step lasted around 3 HRT (7.5 days). Finally, the P.III entitles 120 days in which SMX and TMP were present in the MA-AnMBR feed with a concentration equivalent to 150 µg.L$^{-1}$each.*

### D. Pearson correlation

Pearson correlation tests were conducted between the measured SMX resistant-bacteria, TMP resistant-bacteria, and the different gene relative concentrations, for samples taken from the sludge and the permeate of the MA-AnMBR. A strong correlation was assumed when the absolute value of the Pearson coefficient ($\rho$) was above 0.7. Results of the correlation for sludge and permeate samples can be seen in

**Table D-1** and **Table D-2** respectively.

The sludge SMX resistant bacteria has no lineal correlation with any of the studied genes, but permeate values have a positive linear correlation with the *sul1* gene and *intI-1* MGE (with ρ values of 0.87 and 0.71 respectively). Results of the lack of correlation between SMX-resistant bacteria and genes *sul1* and *sul2* might indicate that the abundance of the selected resistant genes for SMX are not directly linked with the gain of resistance towards the antibiotics expressed in the genes. ARG *sul1, sul2, dfrA1,* and *intI-1* MGE were already present in the MA-AnMBR before the addition of SMX and TMP to the feed. The Ma-AnMBR was inoculated with real sludge from a pilot-scale blackwater anaerobic reactor located at NIOO-KNAW facilities (Wageningen, Netherlands) and therefore, some resistance towards the selected antibiotics could be expected. Thus, the MA-AnMBR could already have some ARGs and the added concentrations of antibiotics might not be enough to assess the gain or loss in resistance.

***Table D-1.*** *Pearson correlation coefficients for sludge samples taken between days 96 and 430 of the MA-AnMBR operation. The genes concentration are considering their relative abundance (over 16S gene abundance), while the resistant bacteria to SMX or TMP correspond to the average plate count on each day. These days correspond to periods during the (stepwise) addition of antibiotics, P.II, after addition, where a concentration of 150 µg.L*$^{-1}$ *of SMX and TMP was added to the MA-AnMBR feed, PIII, and the period entitled P.IV, where SMX and TMP stopped being added to the feed. Strong correlations are shown in bold and with a green background and are defined for absolute values of Pearson coefficients above 0.7.*

| MA-AnMBR sludge | TMP resistant-bacteria | SMX resistant-bacteria | *dfrA1* gene | *sul1* gene | *sul2* gene | *intI-1* MGE |
|---|---|---|---|---|---|---|
| TMP resistant-bacteria | **1.00** | | | | | |
| SMX resistant-bacteria | **0.83** | **1.00** | | | | |
| *dfrA1* gene | -0.35 | -0.23 | **1.00** | | | |
| *sul1* gene | 0.00 | 0.36 | **0.86** | **1.00** | | |
| *sul2* gene | -0.45 | -0.30 | **0.99** | 0.66 | **1.00** | |
| *intI-1* MGE | -0.33 | -0.04 | **0.99** | **0.82** | **0.94** | **1.00** |

***Table D-2.*** *Pearson correlation coefficients for UF permeate samples taken between days 86 and 250 of the MA-AnMBR operation. The genes concentration are considering their relative abundance (over 16S gene abundance), while the resistant bacteria to SMX or TMP correspond to the average plate count on each day. A total of seven sample days were assessed for correlation, from periods P.I (before the addition of antibiotics), P.II (during the stepwise antibiotic addition) and P.III, when the concentration of 150 µg.L*$^{-1}$ *of SMX and TMP was added to the MA-AnMBR feed for around 180 days. Strong correlations are shown in bold and with a green background and are defined for absolute values of Pearson coefficients above 0.7.*

| MA-AnMBR permeate | TMP resistant-bacteria | SMX resistant-bacteria | *sul1* gene | *sul2* gene | *intI-1* MGE |
|---|---|---|---|---|---|
| TMP resistant-bacteria | **1.00** | | | | |
| SMX resistant-bacteria | **0.97** | **1.00** | | | |
| *sul1* gene | **0.94** | **0.87** | **1.00** | | |
| *sul2* gene | -0.17 | -0.11 | -0.22 | **1.00** | |
| *intI-1* MGE | 0.67 | **0.71** | 0.53 | -0.27 | **1.00** |

Mobile genetic element *intI-1* had a positive linear correlation with all tested genes of the MA-AnMBR sludge. Due to their association with plasmids, the class 1 integron plays a key role in the transport of ARGs [1]. The horizontal gene transfer (HGT) linked to *intI-1* concentrations might be responsible for a rise in the extracellular plasmid DNA, which hence intensified the harbouring of plasmid-based resistance within the microorganisms [2]. The relative abundance of the *intI-1* gene in the permeate of the MA-AnMBR was similar to the one obtained in the sludge (Annex C -Gene concentration in the MA-AnMBR**)**. A similar result, where the *intI-1* number of copies in raw and treated sewage had no significant difference was observed by Makowska, et al. [3]. Furthermore, the *sul2* and *intI-1* are found to be co-located on conjugative plasmids generally. The gene cassettes with *sul2* and *intI-1* are found abundantly in wastewater, and the abundance of MGEs like *intI-1* is linked to the presence and distribution of ARG *sul2* [4]. This explains the strong correlation (ρ of 0.94) between *intI-1* and *sul2*. Moreover, the correlation results between *intI-1*, *dfrA1*, and *sul1* might indicate that the MGE *intI-1* could also be located in cassettes with these genes.
